# Live cell imaging and CLEM reveals effects of Mutant Huntingtin Aggregation Process

**DOI:** 10.1101/2023.01.28.526012

**Authors:** Esra Nur Yi̇ği̇t, Mehmet Şerif Aydin, Işıl Aksan Kurnaz, Emrah Eroğlu, Gürkan Öztürk

## Abstract

Huntington’s disease is a neurodegenerative disease caused by expansion of CAG repeats in exon-1 of *Huntingtin (HTT)* gene leading to production of mutant Huntingtin (mHtt) protein. Htt protein is known to play crucial roles in regulation of cytoskeletal dynamics and vesicular transport in physiological conditions. By combining *in vitro* time-lapse imaging and correlative light and electron microscopy (CLEM), we investigated the subcellular dynamics of mHtt during the process aggregate formation. Here we show that distribution of F-actin is affected by mHtt aggregation. F-actin tends to relocate from the peripheral to perinuclear area in cells with mHtt aggregates, possibly caused by sequestration of F-actin from cell membrane to aggregation zones. In accordance with this, mHtt in hippocampal neurons were aggregated into axonal varicosities together with F-actin as they have higher F-actin expression in their neurites compared to their soma. Additionally, correlative light and electron microscopy (CLEM) revealed increased mitochondrial accumulation at the periphery of mHtt aggregates which is surrounded by a F-actin mesh. Mitochondria targeted HyPerRed, a genetically encoded hydrogen peroxide sensor, revealed that oxidative stress in mHtt expressing cells increased. Thus, our findings give new insights about the pathology caused by mHtt through interplay of aggregates with F-actin cytoskeleton and mitochondrial oxidative stress.

## Introduction

Huntington’s disease (HD) is an autosomal dominant neurodegenerative disease caused by a mutation in the coding region of Huntingtin (Htt) protein, resulting with an increased CAG repeats in exon-1 region of huntingtin gene (*Htt*) ^1^. In normal conditions, Htt protein contains less than 36 glutamine residues while pathological cases can have up to 250, and the number of CAG repeats differ between cases and determine the age onset and severity of the disease ^2,3^.

Htt protein is thought to have distinct roles in several physiological processes including neuronal survival, signaling, cellular transport and cytoskeletal dynamics ^4^. Furthermore, it is known that deletion of Htt protein causes lethality in homozygous mice during embryonic period, which implies that it has crucial roles in development ^5^. It is known that expression of full length WT Htt is protective against excitotoxicity caused by NMDA administration ^6^. Immunoprecipitation and immunohistochemistry studies in neurons show that Htt is associated with vesicles, suggesting that Htt may play a role in vesicle trafficking ^7^. One study using Hela cells supported that downregulation of Htt using siRNA reduced the total number of fusion events in the plasma membrane and vesicles travelling from Golgi to ER ^8^.

Expanded glutamine repeats (poly Q) in N-terminus of Htt protein cause gain of toxic function and those Htt proteins are termed as mutant Htt (mHtt). Although it is known that one of the main events happening through the progression of HD pathogenesis is oxidative stress ^9^, the role of mHtt aggregates in this process is not clear yet ^10^. Therefore, it is important to investigate subcellular dynamics through the aggregation of mHtt in high temporal and spatial resolution. To reveal ultrastructural details of mHtt aggregates and their surrounding environment, higher resolution imaging techniques are necessary. In a recent study, correlative light and electron microscopy (CLEM) approach revealed that mHtt aggregates are composed of membranous, fibrillary structures, cytoskeletal proteins and organelle residues ^11^. A study done using super-resolution microscopy showed that polyQ containing aggregate clusters are gathered to perinuclear zone by active transport at the microtubule organization center (MTOC) ^12^.

As Htt is associated with intracellular trafficking in normal conditions, it has been suggested that interaction with ER and cytoskeletal proteins such as microtubules, intermediate filaments and actin filaments are important in pathological conditions. In this context, overexpression of vimentin has been shown to increase mHtt aggregation in Neuro2a cells ^13^, and a study showed that Htt colocalizes to the actin-cofilin rods that are formed in response to heat shock-induced stress ^14^. Additionally, a recent study found that knockdown of Htt and Huntingtin interacting protein 1 (HIP1)- related protein (HIP1R) disrupts SETD2 dependent methylation of actin to some extent and inhibits actin polymerization ^15^. Htt plays a role in regulation of adhesion complexes in fibroblasts by contributing to the accumulation of alpha-actinin, leading to morphological changes in some of the cells ^16^. It was further shown that Htt interacts with beta-tubulin and localizes to the perinuclear area, which enhances its nuclear entry ^17^. These findings support the fact that interaction and interplay of Htt with different cytoskeletal components may have crucial roles in Htt accumulation.

In this study, we transiently expressed mHtt in HEK293T cells and primary hippocampal neurons and demonstrated that mHtt significantly affects actin dynamics resulting in different interaction patterns from coaggregation to nearby accumulation. Ultrastructural analysis revealed mitochondrial accumulation between mHtt aggregates and F-actin mesh. In addition, oxidative stress was increased in accumulated mitochondria which contributes to mitochondrial dysfunction.

## Results

### mHtt in HEK293T cells results with aggregate formation and altered subcellular localization of actin filaments

We expressed mutant (EGFP-Q74) EGFP fusion exon-1 of human Htt protein in HEK293T cells and used wild-type (EGFP-Q23) as its control. We observed that Q74-EGFP Htt forms into aggregates as subcellular bright and condensed EGFP particles while Q23-EGFP Htt shows a diffuse signal pattern (Fig. 1A). Phalloidin staining showed increased cytoplasmic F-actin in Q74-EGFP expressing HEK293T cells compared to Q23-EGFP expression while F-actin levels in cell membrane didn’t change (Fig. 1B, 1D). Further analysis in Q74-EGFP expressing HEK293T cells revealed that cytoplasmic F-actin specifically localized into the periphery of cytoplasmic mHtt aggregates (Fig. 1B, 1D). Co-localization analysis of F-actin with mHtt aggregates didn’t show positive correlation (Pearson correlation coefficient: −0.08), instead it was localized in close proximity to mHtt aggregates (Fig. 1B). HEK293T cells were co-transfected with LifeAct-tdTom and Q23-EGFP or Q74-EGFP for F-actin imaging in living cells. LifeAct-tdTom showed similar F-actin pattern compared to phalloidin staining, cytoplasmic F-actin is concentrated near mHtt aggregates (Fig. 1C).

**Figure 1:**
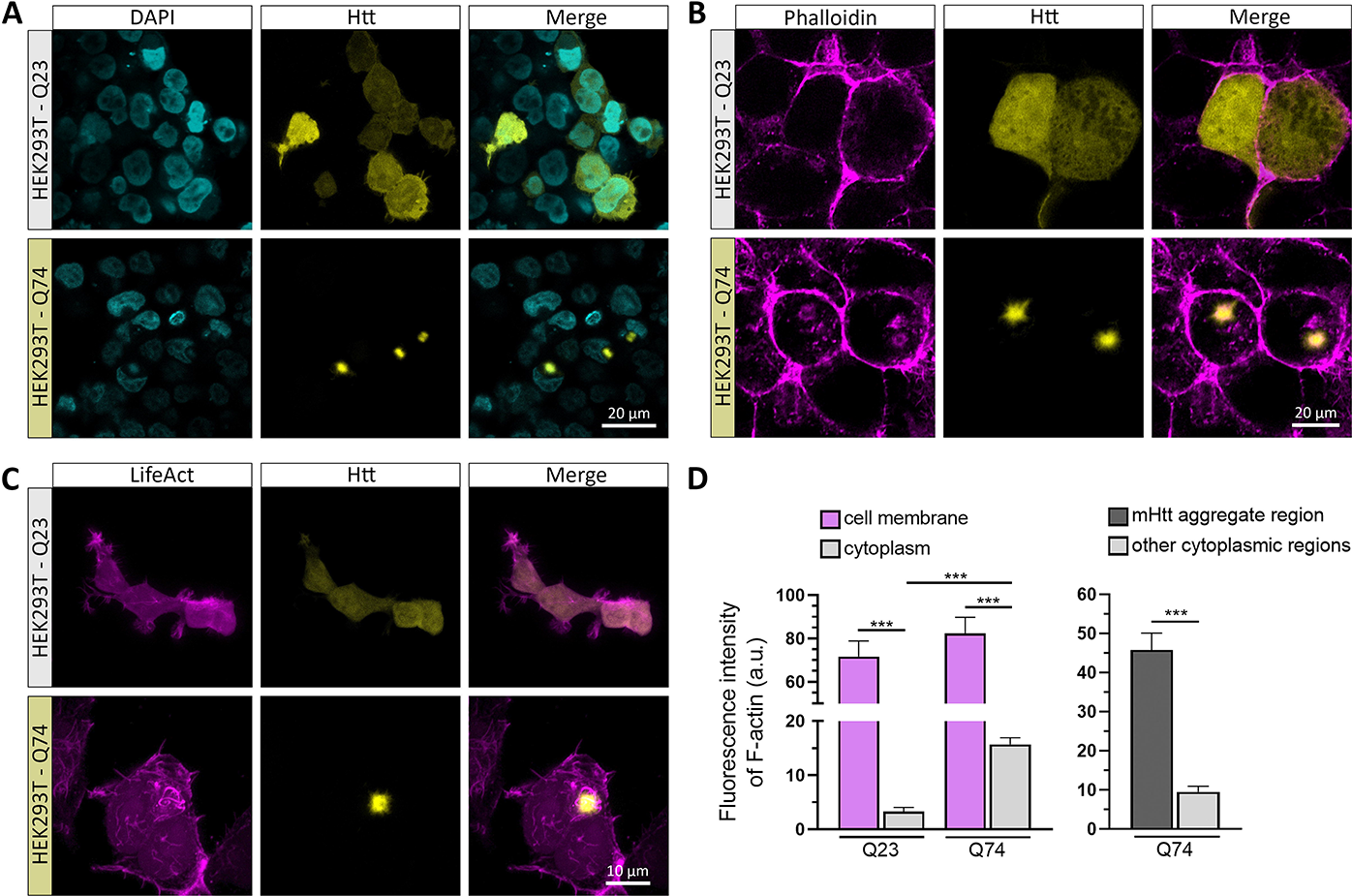
Co-localization of F-actin with mHtt aggregates in HEK293T cells. (A) Representative confocal microscopy images of Q23-EGFP (wild-type Htt) and Q74-EGFP (mutant Htt) expressing (yellow) HEK293T cells. mHtt aggregates appeared as dense and bright particles (lower panel). Nucleus were counterstained with DAPI (turquoise). (B) Confocal microscopy images of Q23-EGFP and Q74-EGFP expressing (yellow) HEK293T cells stained with phalloidin (magenta). (C) Confocal microscopy images of LifeAct-tdTom (magenta) expressing HEK293T cells co-transfected with either Q23-EGFP or Q74-EGFP (yellow). (D) Quantification of fluorescence intensity of F-actin in membrane and cytoplasm of phalloidin stained HEK293T cells expressing Q23-EGFP and Q74-EGFP (left panel, n=100, Student’s t test was performed for statistical analysis). F-actin accumulation by fluorescence intensity in Q74-EGFP expressing HEK293T cells. F-actin expressions were measured in mHtt aggregate region and other regions in the cytoplasm of same cell (right panel, n=20, Student’s t test was performed for statistical analysis.). Error bars in graphs are S.E.M.

### F-actin co-accumulates with mHtt aggregates in HEK293T cells

We co-transfected HEK293T cells with LifeAct-tdTom and Q23-EGFP or Q74-EGFP and performed time lapse confocal microscopy imaging after 24h of transfection. F-actin was localized into cell membrane in Q23-EGFP expressing HEK293T cells while it was located near mHtt aggregates in Q74-EGFP expressing HEK293T cells through the aggregation process (Fig 2A). High resolution Z-stack confocal microscopy images of HEK293T cells were taken after time lapse imaging and same cells were prepared for array tomography. Serial transmission electron microscopy (TEM) images correlated with confocal images enabled visualization of mHtt aggregates in HEK293T cells at high resolution. In fact, even in TEM images only without any labeling the aggregation regions could be spotted as electron dense areas (Fig. 2D_1_ – inset). CLEM images showed that membranous and fibrillary structures were entangled within mHtt aggregates, exhibiting a complex and irregular architecture. 3D reconstruction revealed excessive mitochondrial accumulation (red) and increased ER network around mHtt aggregates (Fig. 2C, 2D). Mitochondria were accumulated between F-actin mesh and mHtt aggregates (Fig. 2C).

**Figure 2:**
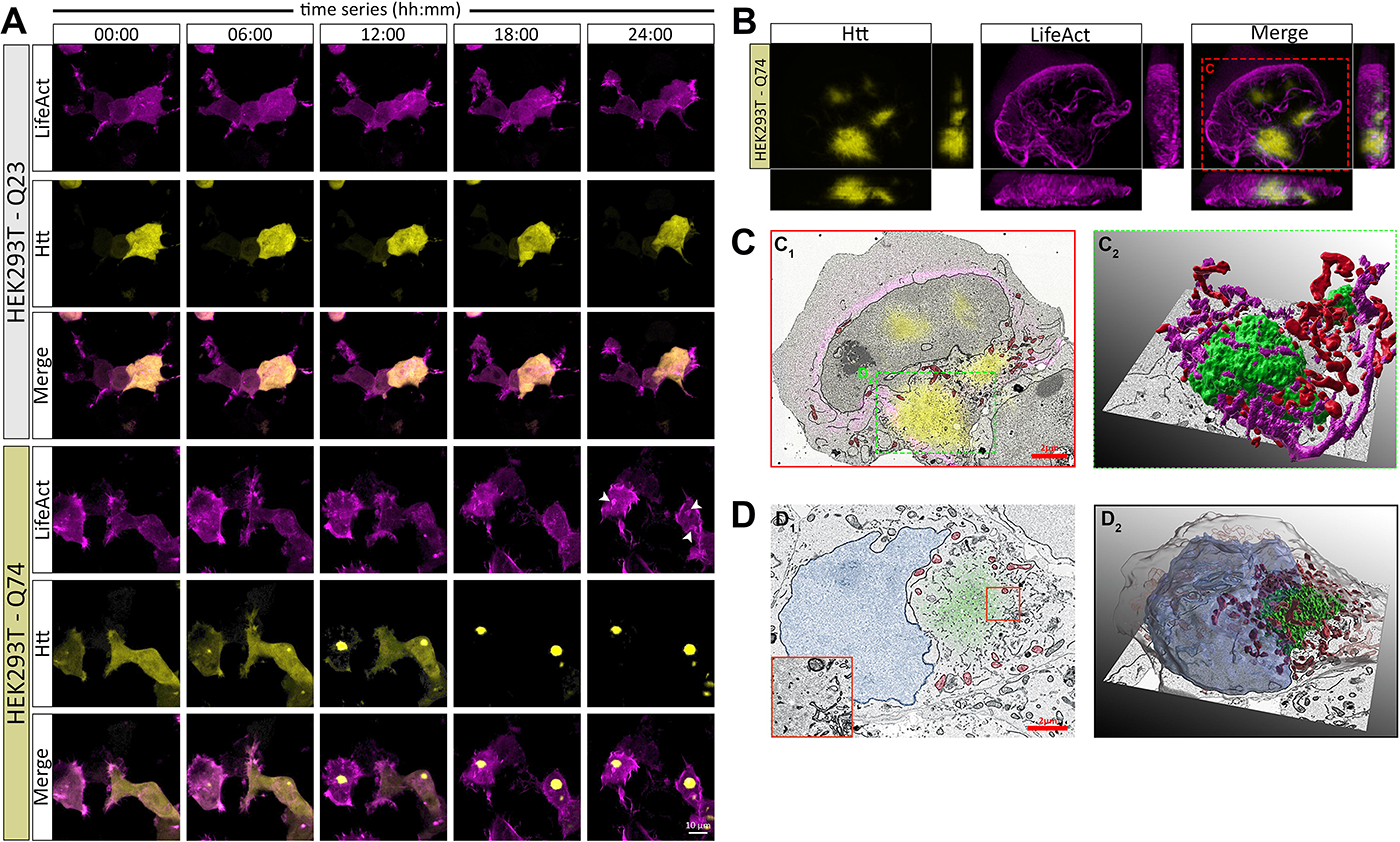
mHtt dependent F-actin and mitochondrial accumulation in HEK293T cells. (A) Timelapse confocal microscopy imaging of F-actin accumulation and mHtt aggregation in LifeAct-tdTom with Q23-EGFP and LifeAct-tdTom with Q74-EGFP expressing HEK293T cells. White arrows indicate accumulation of F-actin with mHtt aggregates. (B), (C) CLEM image of LifeAct-tdTom and Q74-EGFP expressing HEK293T cell. (B) Confocal microscopy image of nuclear and cytoplasmic mHtt aggregates (yellow) and F-actin (magenta). (C1) TEM image of LifeAct-tdTom and Q74-EGFP expressing HEK293T cell indicating mHtt aggregates (yellow), F-actin (magenta) and mitochondria (red). (C2) 3D reconstruction of TEM images of serially sectioned cytoplasmic mHtt aggregate (green), F-actin (magenta) and mitochondria (red). (D1) TEM image of Q74-EGFP expressing HEK293T cell indicating mHtt aggregate (green) and mitochondria (red). (D2) 3D reconstruction of TEM images of serially sectioned Q74-EGFP expressing HEK293T cell showing the outer border of cell (gray), nucleus (light blue), perinuclear localized mHtt aggregate (green), and mitochondrial accumulation (red) around the mHtt aggregate.

### Oxidative stress was increased in mitochondria of Q74-EGFP expressing HEK293T cells

In order to investigate if there is a relation between mitochondrial accumulation and oxidative stress in HEK293T cells mHtt aggregates, we co-transfected HEK293T cells with a red fluorescent hydrogen peroxide sensor HyPerRed-mito and Q23-EGFP or Q74-EGFP (Fig. 3A). Time-lapse confocal microscopy imaging after 24h of transfection showed a gradual increase in mitochondrial HyPerRed fluorescent intensity in Q74-EGFP expressing HEK293T cells compared to wild-type control Q23-EGFP (Fig. 3B-left panel). Fluorescent intensity analysis at the end of time lapse imaging (24h) showed this increase was significantly higher in Q74-EGFP expressing HEK293T cells compared to Q23-EGFP expressing HEK293T cells (Fig 3B-right panel). Additionally, it is found that fluorescent intensity of mitochondrial HyPerRed is significantly higher in Q74-EGFP expressing HEK293T cells which have mHtt aggregates compared to the ones that do not have (Fig 3B).

**Figure 3:**
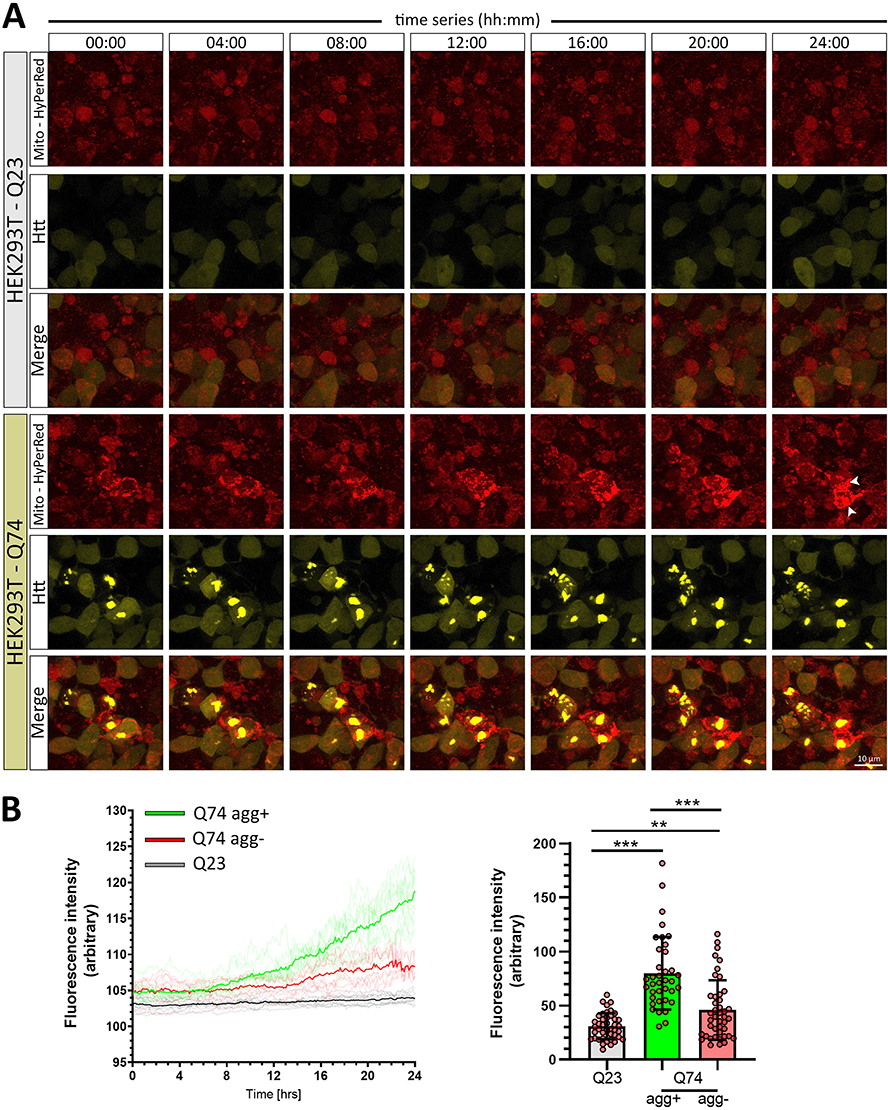
Mitochondrial oxidative stress is increased in mHtt expressing HEK293T cells. (A) Time-lapse confocal microscopy images of HyPerRed-mito (red) and Q23-EGFP or Q74-EGFP (yellow) expressing HEK293T cells. (B) Fluorescence intensity quantification of mitochondrial HyPerRed in time-lapse recordings of Q23-EGFP expressing and Q74-EGFP expressing HEK293T cells with (agg+) or without (agg-) aggregated mHtt (left panel, n=10). Fluorescence intensity quantification and statistical analysis of mitochondrial HyPerRed at the end of time-lapse recordings of Q23-EGFP expressing and Q74-EGFP expressing HEK293T cells with or without aggregated mHtt (right panel, n=42, 37 and 43 respectively, Student’s t test was performed for statistical analysis).

### F-actin co-accumulates in axons of Q74-EGFP expressing neurons together with mitochondria accumulation

In order to investigate mHtt aggregation process in neurons, hippocampal neurons were co-transfected with LifeAct-tdTom and Q23-EGFP or Q74-EGFP. Similar to the findings in HEK293T cells, aggregated mHtt were appeared as bright and dense EGFP particles while wildtype Htt showed a diffuse pattern (Fig 4A). To examine the relation between axonal mHtt and F-actin in detail, we performed time-lapse tracking analysis; we showed that mHtt aggregates were smaller at the beginning of the aggregation process (Fig. 4B - see time points 00:00, 01:00 and 02:00, Supplementary movie 3). As smaller fragments are gathered and accumulated together, which can be observed as increased fluorescence intensity, their motility was found to decrease (Fig. 4B 07:00 onwards, see right panel for velocity quantification). Interestingly, as motility of axonal mHtt aggregates decreased and they grew larger in size, varicosities were observed in neurites (Fig. 4C – upper panel). Time-lapse tracking revealed that the larger the mHtt aggregates got, the more F-actin was accumulated at these sites (Fig. 4C).

**Figure 4:**
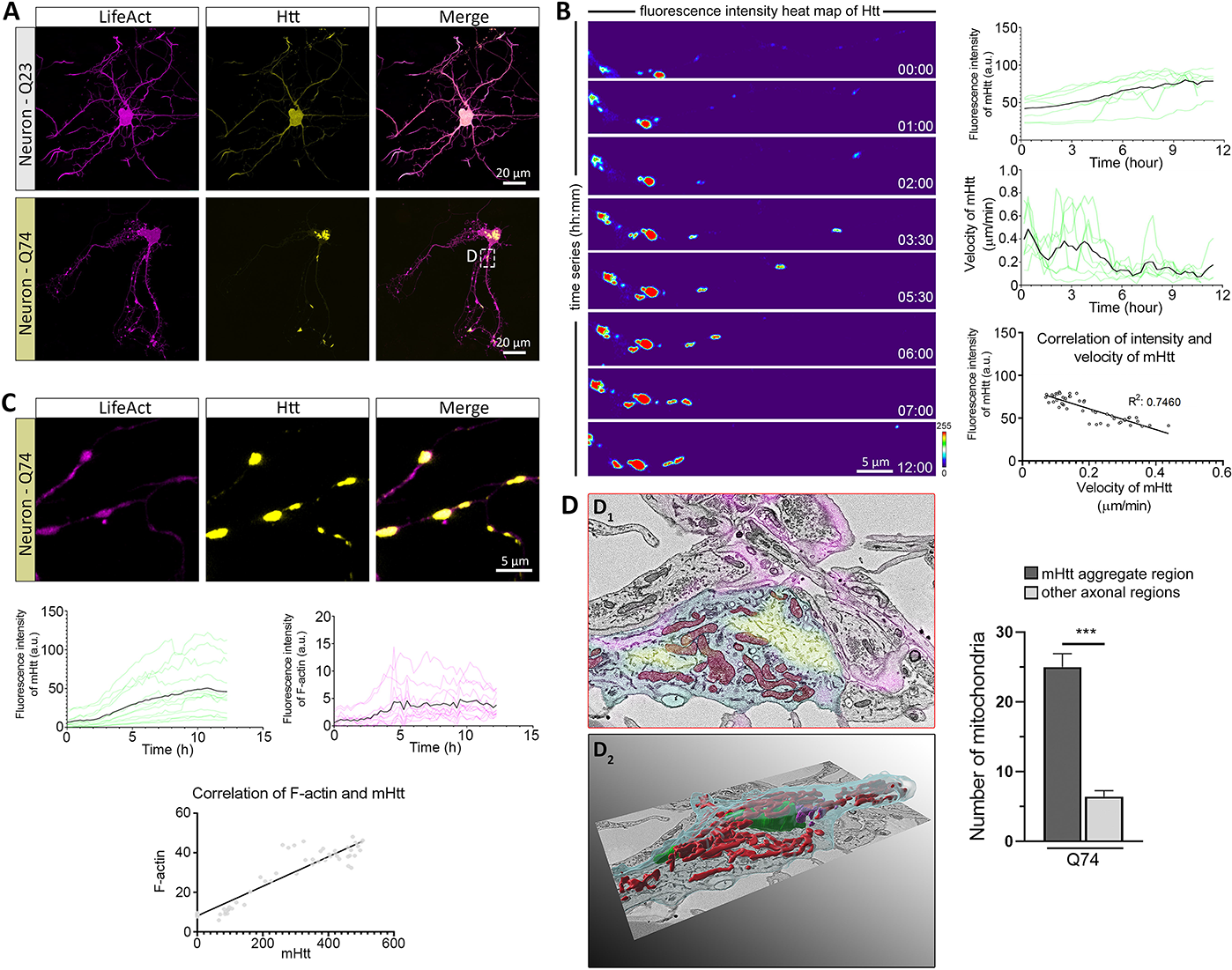
Aggregation of axonal mHtt and co-accumulation of axonal F-actin and mitochondria in neurons. (A) Representative confocal microscopy images of LifeAct-tdTom and Q23-EGFP (wild-type Htt) or Q74-EGFP (mutant Htt) expressing (yellow) neurons. Cytoplasmic and axonal mHtt aggregates appeared as dense and bright fluorescent particles (lower panel). (B) Retrograde transport of axonal mHtt aggregates indicated by fluorescence intensity heat map of timelapse confocal imaging in hippocampal neurons. Soma of the cell resides at the leftmost side (left panel). Velocity and fluorescence intensity values of axonal mHtt aggregates were quantified by time-lapse tracking (n=7). Correlation of fluorescence intensity and velocities of mHtt aggregates (bottom panel) during time lapse imaging. (C) Axonal mHtt aggregates (yellow) accumulated into F-actin (magenta) rich neurite varicosities in Q74-EGFP and LifeAct transfected hippocampal neurons (upper panel). Fluorescence intensity of mHtt (green) and F-actin (pink) tracked over time and normalized against background signal in axons (n=13). Correlation of fluorescence intensity and velocities of F-actin rich zones (bottom panel) during time-lapse imaging. (D) TEM image (D1, yellow: mHtt aggregate, red: mitochondria, magenta: F-actin, blue: neurite) and reconstruction (D2, green: mHtt aggregate, red: mitochondria, magenta: F-actin, blue: neurite) of axonal mHtt aggregate with mitochondrial accumulation. Quantification of axonal mitochondria number accumulated into the mHtt aggregate region in Q74-EGFP expressing hippocampal neuron compared to that of other axonal regions containing no mHtt aggregates.

Correlation of confocal and TEM images with CLEM technique revealed that actin filaments surrounding mHtt aggregates formed a mesh-like structure that contains high number of mitochondria in neurites (Fig. 4D). CLEM image showed that F-actin accumulation was primarily found in axonal mHtt aggregates and mitochondrial accumulation was prominent in those regions (Fig. 4D_1_). Quantification of axonal mitochondria proved that number of mitochondria was significantly increased near axonal mHtt aggregates compared to other axonal regions (Fig. 4D_2_).

## Discussion

Huntingtin is a multifunctional protein that takes place in various cellular processes almost all of which is related to cytoskeleton and its main components tubulin and actin ^4^. Compared to other cells, neurons have a more elaborate cytoskeleton extending into dendrites and axons, which can be longer than 1 m. For proper functioning and maintenance of a neuron, transport of molecules and organelles between soma and neurites should be efficiently maintained throughout its life. It is not surprise that in most neurodegenerative diseases, this transport is defective though it is still unclear whether this precedes or follows somatic ailments^18^. In this study, we exploited live cell imaging to better understand the interplay between mHtt, F-actin and mitochondria. Combining high resolution correlative microscopy, time-lapse imaging and genetically labelled proteins and biosensors we analyzed F-actin and mitochondria accumulation together with mHtt aggregation in HEK293T cells and hippocampal neurons.

In HEK293T cells transfected with mHtt, the aggregates were found to predominantly appear in the perinuclear region as a solitary dense inclusion and less in the nucleus (data not shown). This distribution pattern supports the suggestion that since Htt is closely related to tubulin and takes part in motor transport, it tends to aggregate over tubulin network which is richest at the centrosome typically located in the perinuclear zone ^4,17^. Indeed, in neurons, which do not use centrosomes to nucleate tubulin ^19,20^, mHtt aggregation occurred irregularly as fragmented and less dense cytoplasmic and more prominent nuclear inclusions. It is reported that Htt mRNA is concentrated in the nucleus of neurons, which may facilitate the perinuclear translation of the protein and its transport into the nucleus ^21^. Consistent with other reports ^22^ we observed accumulation of mitochondria around the mHtt aggregates in HEK293T cells, which were probably entrapped by dense mHtt aggregates.

Experiments with LifeAct, a special peptide that fluorescently labels actin without interfering with its dynamics ^23^, revealed details about interaction of actin and mHtt. In HEK293T cells, we consistently observed that actin is accumulated around the perinuclear zone and it surrounds periphery of mHtt aggregates. This distribution pattern, which is also reported by others ^22^, couldn’t observed in hippocampal neurons. To explain this difference, we developed a hypothesis suggesting that actin is sequestered by mHtt. Support to this first comes from the literature suggesting that Htt may bind to actinin, a protein that bundles and crosslinks actin filaments and huntingtin interacting protein (HIP) ^16,24^. We demonstrated that HEK293T cells have relatively more actin in their cytoplasm and at the peripheral zones compared to hippocampal neurons (Supplementary Fig. 1). We suggest that above a certain concentration F-actin may bind to mHtt and co-transported towards perinuclear region or even co-aggregate. Another important observation was that in HEK293T cells bulks of actin were moved from the periphery towards nucleus and stuck to mHtt aggregate (Supplementary Fig. 2). Displacement of F-actin from different parts of the cell to aggregation zones may be related to impaired Htt functions. Wild type Htt was previously shown to recruit actinin to focal adhesions and contributes to formation of stress fibers and adhesion complexes ^16^. It was further shown that as aggregates form, mHtt sequesters wild type Htt ^25,26^ resulting in impaired Htt activity which may help actin bundles be dislodged from these sites and hooked and dragged by mHtt centrifugally.

Oxidative stress is known to involve in the pathogenesis of HD but it remains to be clarified if it is an indirect outcome of pathogenesis or have a direct relationship with the Htt aggregate formation ^9,10,27^. Utilization of mitochondria localized HyPerRed, a genetically encoded hydrogen peroxide biosensor ^28^, have provided important findings about the relation of oxidative stress and mHtt aggregation process. It is found that expression of mHtt was sufficient for increased levels of mitochondrial hydrogen peroxide compared to expression of wild-type Htt. Besides Q74-EGFP expressing cells with visible mHtt aggregates showed significantly higher hydrogen peroxide levels compared to the ones that do not yet have aggregates. These findings are in accordance with research that show oxidative stress is related to the mHtt aggregation ^29^. However, the cause-and-effect relationship between mHtt aggregate formation and mitochondrial oxidative stress is yet to be consolidated.

An important finding of this study was that in the neurites of hippocampal neurons, co-aggregation of F-actin and mHtt causes formation of varicosities, an early change prior to axon degeneration, most likely by disrupting axonal transport ^30,31^. Consistent with our findings, others reported that axonal mHtt aggregation precedes somatic changes in animal models of Huntington ^32–34^, which suggests that Huntington’s disease may start as an axonopathy ^35,36^. Further research with neurons is also necessary to determine the oxidative stress and its relation with axonal transport deficit in more detail.

In conclusion, in this study we showed that while aggregating, mHtt closely interacts with F-actin and this results in its co-aggregation or accumulation around mHtt aggregates. Localization of mitochondria with increased oxidative stress around these aggregation zones also suggests impairment of mitochondrial transport, which will further contribute to neurodegeneration.

---

**Supplementary Figure 1:**
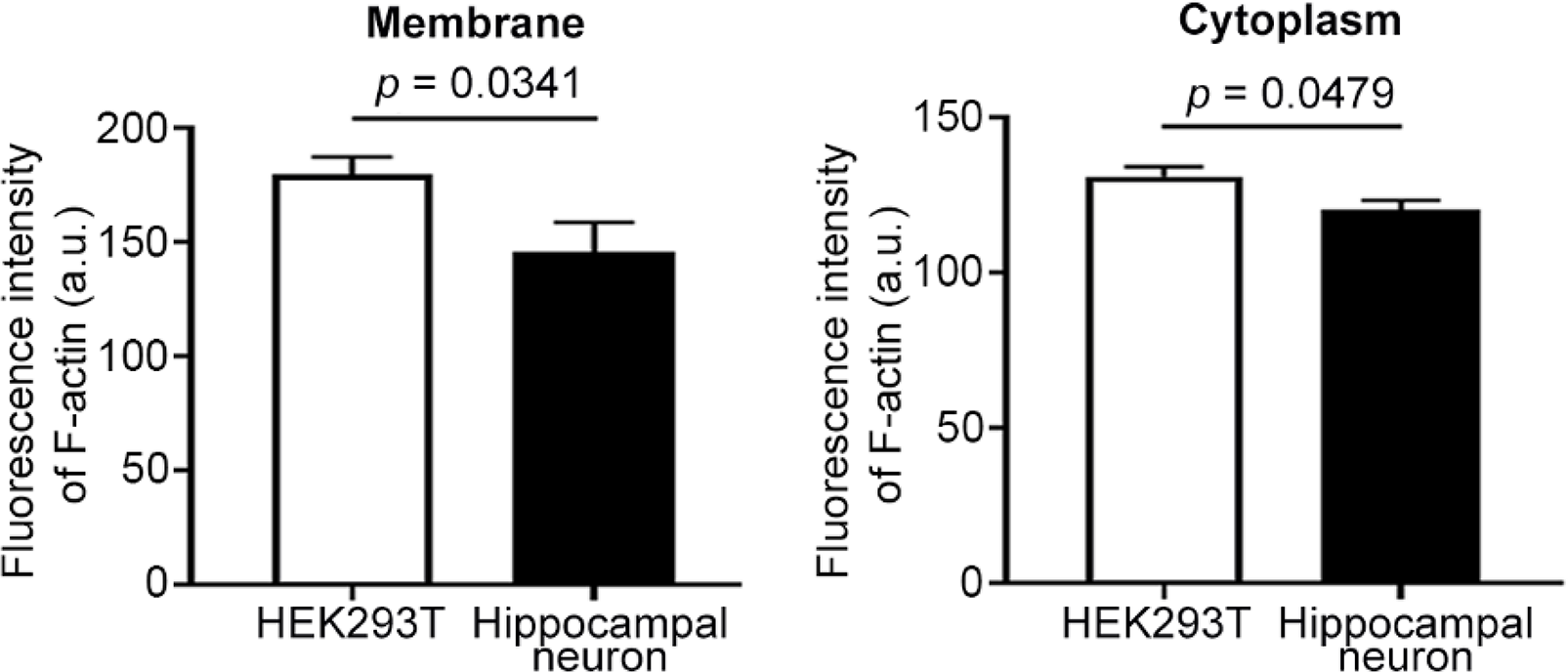
F-actin distribution in phalloidin stained HEK293T cells and neurons.

**Supplementary Figure 2:**
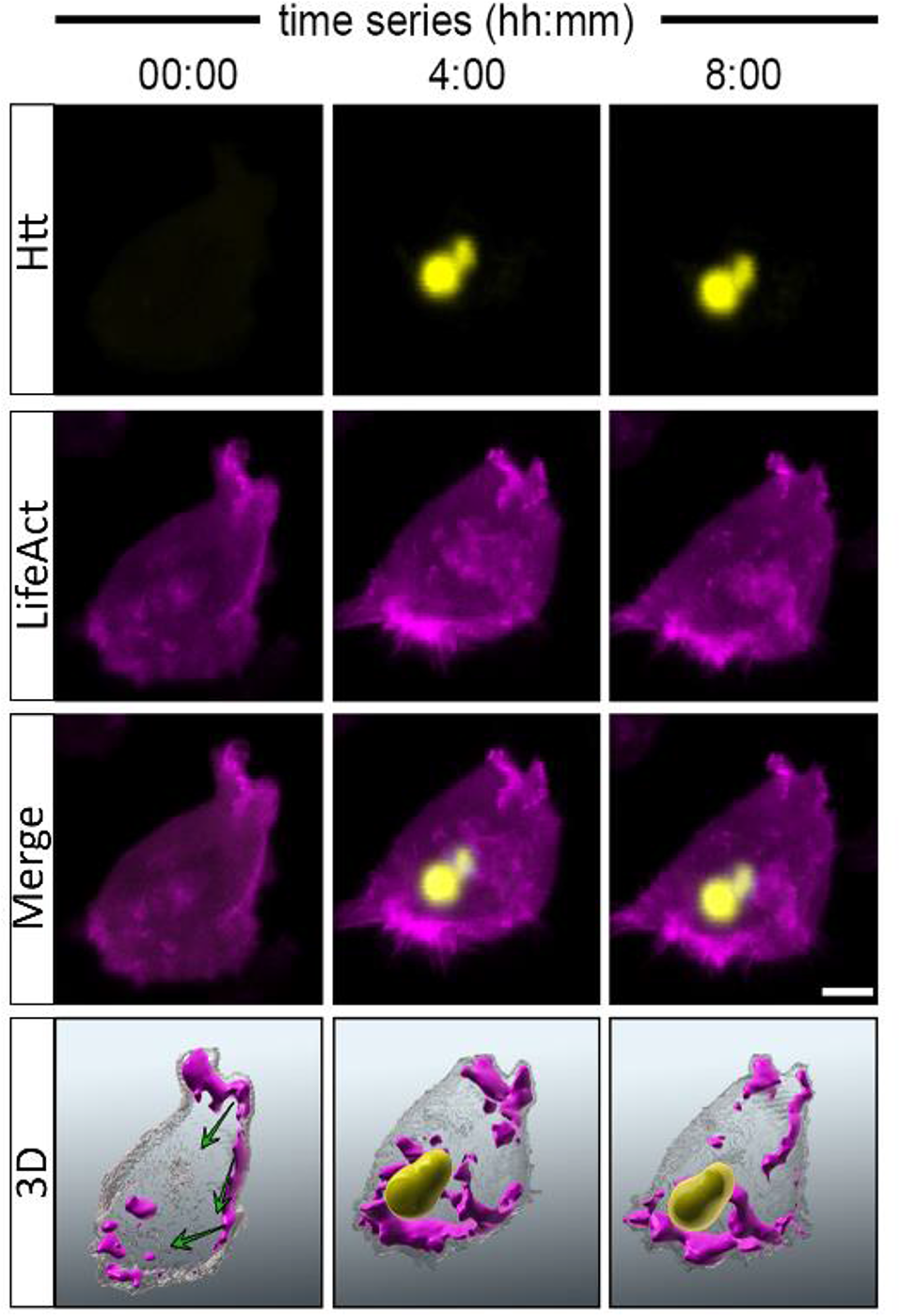
Time-lapse confocal microscopy images representing the relocation of F-actin in Q74-EGFP and LifeAct expressing HEK293T cells. Green arrows indicate movement pattern of f-actin from membrane to mHtt aggregate region through the time.

## Materials and Methods

### Cell culture, Transfection and Live Cell Imaging

HEK293T cells were maintained in high glucose DMEM (41966-029, Gibco) supplemented with 10 % FBS (10500-064, Gibco) and 1 % solution (450-115-EL, MultiCell). Cells were plated on 100 mm cell culture dishes or poly-D-lysine coated glass bottomed 8-well plates on the day before transfection for western blot and correlative light electron microscopy (CLEM) respectively. Expression plasmids were transfected using polyethylenimine (PEI) (23966-1, Polysciences). 10 ug and 200 ng of total plasmid was used for 100 mm cell culture dishes and 8-well plates respectively with a Plasmid:PEI ratio of 1:4. Briefly, plasmid:PEI mixtures were prepared in serum-free DMEM and vortexed for x15 pulses. After 15-20 min incubation at room temperature, plasmid:PEI mixtures were added to cells and fresh medium was added after 16h. pEGFP-Q23 and pEGFP-Q74 were gifts from David Rubinsztein (Addgene plasmid # 40262; http://n2t.net/addgene:40262; RRID:Addgene_40262). pLenti-LifeAct-tdTomato was a gift from Weiping Han (Addgene plasmid # 64048; http://n2t.net/addgene:64048; RRID:Addgene_64048). HyPerRed-mito was a gift from Vsevolod Belousov (Addgene plasmid # 60247; http://n2t.net/addgene:60247; RRID:Addgene_60247) Q23-EGFP or Q74-EGFP expression plasmids were co-transfected with Lifeact-tdTomato or HyPerRed-mito.

For live cell imaging, time lapse recordings were taken with confocal microscope (Carl Zeiss, LSM 880) using Plan-Apochromat 40x/1.3 oil immersion objective in an incubation chamber with 20 min time interval for HEK cells and 15 min time interval for neurons. All experimentation was started 24h after transfection of pEGFP-Q74 unless otherwise specified.

### Primary Neuron Culture

Animal experimentation was done with the approval of İstanbul Medipol University Animal Experimentation Local Ethical Committee. Primary hippocampal cells were isolated from neonatal mice and cultured separately. Briefly, postnatal 1-3 days mice were euthanized and brains were collected under aseptic conditions. Hippocampus were dissected carefully in dissection medium (L15, 1% antibiotic-antimycotic solution), cut into small pieces and put into separate petri dishes. Hippocampus were digested with papain (Sigma, 12.5 U/ml and 25 U/ml respectively) for 45 min with agitation at 4 °C. Afterwards, tissues were triturated via serial pipetting and papain is inhibited with 10 % FBS containing dissection medium for 15 min at 4 °C. Then, cells were pelleted by centrifugation at 180 g for 5 min at 4 °C, dissection medium was discarded and cells were plated into poly-D-lysine coated 8-well glass bottomed dishes. Cells were co-transfected with Q74-EGFP and LifeAct-tdTomato using lipofectamine at a ratio of 1:3 at div 10 and imaged at 12^th^ h of transfection.

### Correlative Light Electron Microscopy (CLEM)

Sample preparation protocol for correlative microscopy was adapted from Ashaber et al.^37^. Cells were washed with 0.2 M sodium cacodylate buffer and fixed using Karnovsky’s fixative (2% paraformaldehyde, 2.5% glutaraldehyde in 0.15 M cacodylate buffer containing 2 mM CaCl_2_) for overnight at +4 C followed by three sequential washes with ddH_2_O. Regions were selected for CLEM and airy scan and/or confocal images were taken using confocal microscope (Carl Zeiss, LSM 880) using Plan-Apochromat 40x/1.3 oil immersion objective with 488 nm and 541 nm excitation lasers. With a 355 nm UV Laser (Rapp Optoelectronic) regions of interests (ROI) were marked by etching glass coverslips.

Cells were post-fixed in reduced OsO_4_ (2% OsO_4_ + 1.5 potassium ferrocyanide) for 30 min at room temperature. All subsequent washes were done using ddH2O unless otherwise specified. Fixed cells were washed for three times and incubated in 0.5 % Thiocarbohydrazide for 15 min at room temperature. After three sequential washes, cells were incubated with 2% OsO_4_ for 30 min at room temperature and washed for three times again. Then, cells were incubated in 2% Uranyl Acetate overnight at +4°C and washed three times. Lead aspartate was added to the cells and incubated at 60°C for 25 min. Afterwards, cells were washed five times thoroughly and dehydrated with graded EtOH series (50%, 70%, 90 %, 100 %, 100 %) 5 min each step. Finally, cells were incubated in 1:1 and 2:1 Epon:EtOH mix respectively, embedded in 100% epoxy resin (Sigma Aldrich, 45359) and incubated at 60 °C until polymerized.

### Array Tomography

To mark ROIs in polymerized blocks, laser etchings on cover glass were localized using Plan-Apochromat 10x/0.4 objective and laser etched on the epoxy blocks. Cover glasses were removed from epoxy blocks using sequential liquid nitrogen/heat bath incubation. Epoxy blocks were trimmed according to the ROI marks and 60 nm serial sections were taken using ultramicrotome (Leica EM UC7) with a diamond knife (Diatome). Cover glasses were manually cut into small pieces and fire-polished. Serial sections were collected onto cover glasses and coated with gold with sputter current 30 mA for 5 seconds (Leica EM ACE200). Cover glasses were sealed on aluminum stabs using carbon tape. Electron microscopy images were taken with Zeiss Gemini SEM 500 using 3.00 kV accelerating voltage and backscatter electron detector with 6.00 K X magnification and 4096x3072 pixel resolution.

### Image Processing & Segmentation

For fluorescence intensity analysis and aggregate tracking, Fiji ^38^ and ZEN software (Carl Zeiss Microscopy GmbH, Germany) were used. For TEM images, serial sections were automatically aligned using Fiji and fine adjustments were done with Adobe Photoshop. Aligned serial TEM and confocal images were segmented with Ilastik ^39^. Both images and segmentations were correlated using Fiji, Photoshop and ZEN software.

### Statistics

The Graph Pad Prism 6 (GraphPad Software Inc., CA, USA) was used for statistical analysis. Student’s t test, and Pearson’s correlation analyses were used where appropriate.

## Acknowledgments

This project was financially supported by “The Scientific Research Projects (BAP) Council of Gebze Technical University” with a grant number with project no: 2018-A101-03.

## Author Contributions

E.N.Y and G.O. wrote the manuscript, E.N.Y., G.O., E.E. and I.A.K. designed experiments, review the manuscript, E.N.Y., and M.S.A. perform the experiments, analyzed data, interpreted results.

