## Supplementary figures and images for "Live cell imaging and CLEM reveals effects of Mutant Huntingtin Aggregation Process"

### Supplemental Figure 1

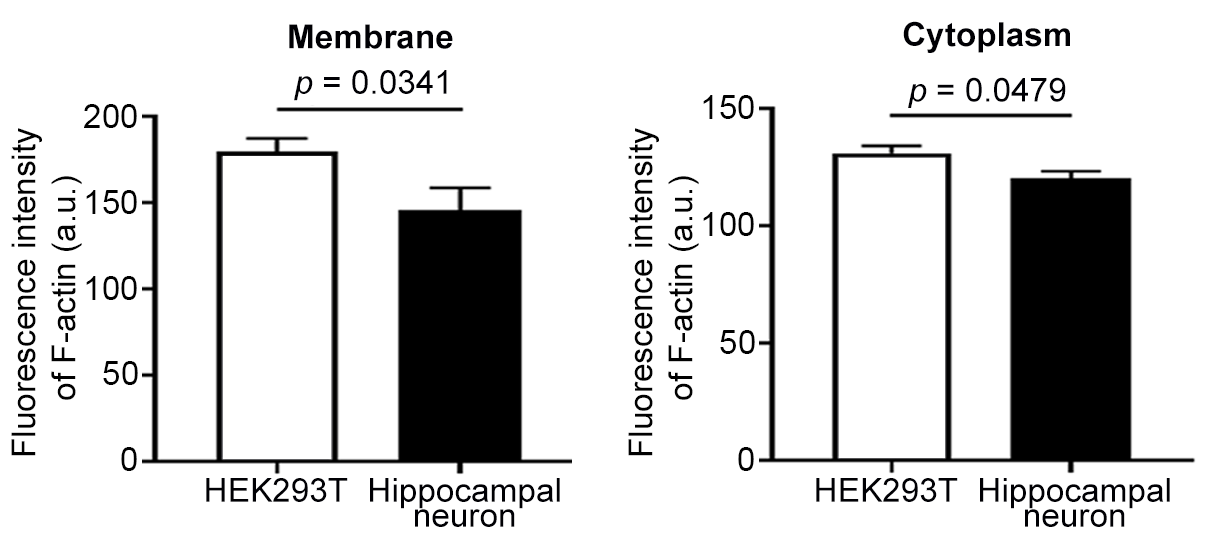

### Supplemental Figure 2

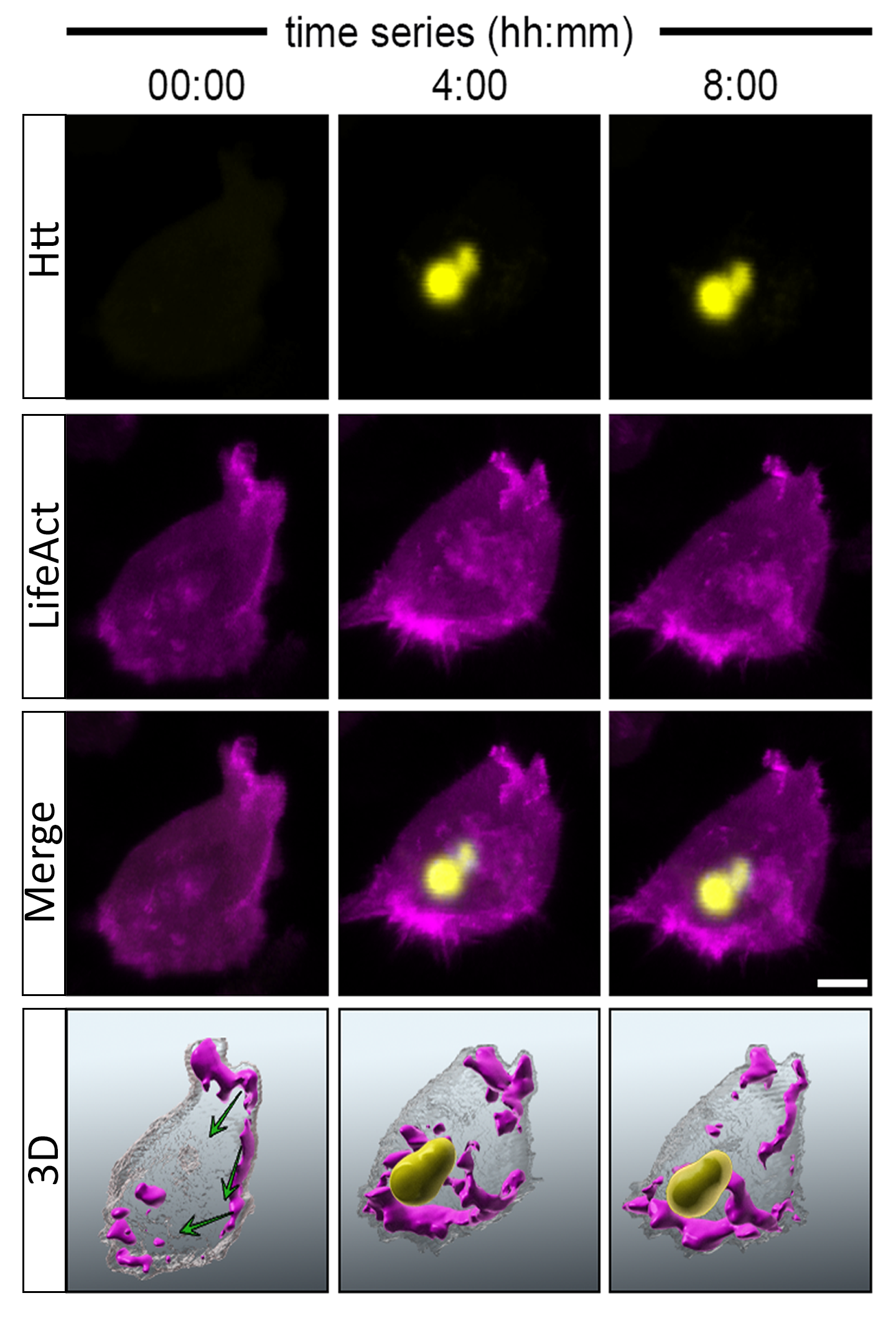
